# Excessive apoptosis of naïve T cells resulting hyperactivation as a cause of mammalian aging

**DOI:** 10.1101/2023.02.24.529806

**Authors:** Lingxia Wang, Xixi Zhang, Haiwei Zhang, Kaili Lu, Ming Li, Xiaoming Li, Yangjing Ou, Xiaoming Zhao, Xiaoxia Wu, Xuanhui Wu, Jianling Liu, Mingyan Xing, Han Liu, Yue Zhang, Yongchang Tan, Fang Li, Jiangshan Deng, Xiaojie Zhang, Jinbao Li, Yuwu Zhao, Xiuzhe Wang, Yan Luo, Ben Zhou, Haibing Zhang

## Abstract

In mammals, the most remarkable T cell variations with aging are the shrinking of naïve T cell pool and enlargement of memory T cell pool, which are partially caused by thymic involution. However, it remains an enigma whether these T-cell-related changes are consequences or causes of mammalian aging. In this study, we find that the T-cell specific *Rip1* KO mice present similar age-related T cell changes and exhibit signs of accelerated aging, including inflammation, multiple age-related diseases and a shorter lifespan. Mechanistically, T cells lacking RIP1 displayed excessive apoptosis, leading to T cell compensatory proliferation, hyperactivation, increased inflammation, and ultimately premature death. Consistent with this, blocking apoptosis by co-deletion of *Fadd* in *Rip1* deficient T cells significantly recovered the lymphopenia and imbalance between naïve and memory T cell, substantially restored ageing-related phenotypes, and prolonged life span in T-cell specific *Rip1* KO mice. These results suggest that changes in T cells play a causal role in mammalian aging. Therefore, replenishing or blocking apoptosis of naïve T cells could offer new therapeutic approaches for aging and age-related diseases.

## Introduction

Human aging is characterized by a gradual decline in the physiological and functional capacity of tissues and organs, resulting in multimorbidity and mortality^1^. Aging is a major risk factor for the development of many chronic diseases, including cancer, cardiovascular disease, neurodegeneration, and infectious diseases^2^. Research has shown that aging disrupts immune functions over time, making individuals more susceptible to various diseases^3^. On the other hand, accumulating evidence shows that changes in the immune system can also contribute to the onset of aging and accelerate the aging process^4^. Thus, the relationship between aging and the immune system remains highly complex and intricate, and much of it is still not fully understood.

The role of the innate immune system in age-related health problems has been extensively studied^4^. However, recent studies have also revealed the active involvement of adaptive immune system in such processes, with a focus on T cells^5^. Exhausted T cells have been found to produce high levels of granzyme K, exacerbating inflammation and suggesting that different subsets of age-related T cells may promote tissue damage^6^. External factors such as chronic viral infections can accelerate the accumulation of these age-related T cells^7^. Additionally, T cell-specific deletion of the mitochondrial transcription factor A (TFAM) not only causes immunometabolic dysfunction that drives T cell senescence, but also leads to a general decline in overall health and the emergence of multiple aging-related features^8^, highlighting the crucial role of T cell aging in overall deterioration.

It is now well recognized that multiple age-related biological changes can lead to defective T cell responses, but the mechanisms by which these changes affect T cells are not yet clear^5^. One of the most striking features of aging is thymic involution, in which the thymus degenerates over time and reduces the replenishment of naive T cell pools necessary for protective immunity^9–12^. Impaired naive T cell output disrupts the composition of peripheral T cells, resulting in compensatory expansion of pre-existing T cell clones and hyperactivation. These adaptations lead to decreased TCR diversity, impaired TCR responses and altered distribution of T cell subsets^13^. Such cellular changes and clonal expansion may contribute to the loss of immune responses to diverse antigens in older adults. However, whether these T cells related changes are a consequence or a cause of mammalian aging remains an enigma.

Receptor-interacting protein kinase 1 (RIPK1 or RIP1) plays a crucial role in regulating cell survival, inflammation, and either apoptosis or necroptosis, depending on the cell context^14^. Previous research found that *Rip1*-deficient mice die at birth from systemic inflammation, which can be prevented by blocking both FADD/Caspase8-dependent apoptosis and RIP3/MLKL-dependent necroptosis^15, 16^. Patients with RIP1 deficiency experience recurrent infections, early-onset inflammatory bowel disease, and progressive polyarthritis^17–19^. More importantly, they exhibit T-lymphopenia, which is consistent with the phenotype of mice with T cell–specific deletion of *Rip1*, due to excessive apoptosis^20^. However, it is still unknown whether the pronounced T-lymphopenia and hyperactivation in T cell–specific *Rip1*-KO mice could cause aging-related diseases or premature death.

In this study, we observed that mice lacking *Rip1* in T cells (*Rip1^fl/fl^Cd4^Cre^* mice, called *Rip1^tKO^* hereafter) showed signs of accelerated aging, which eventually resulted in premature death. Mechanistically, T cells with *Rip1* deficiency underwent excessive apoptosis, leading to compensatory proliferation and hyperactivation, as well as increased levels of inflammation, which eventually caused a shorter lifespan. Importantly, blocking necroptosis had little effect on the T cell compartment or alleviating the multimorbidity and premature aging seen in *Rip1^tKO^*mice. In contrast, blocking apoptosis by deleting *Fadd* in *Rip1* deficient T cells significantly improved the T-lymphopenia, restored the balance between naïve and memory T cells, rescued the aging-related phenotypes and prolonged the shortened lifespan. These results suggest that the apoptosis of naïve T cells, which leads to compensatory proliferation and an increase in memory T cells, could be a cause of aging in mammals. Therefore, replenishing or inhibiting apoptosis of naïve T cells could be a novel approach for treating aging and age-related diseases.

## Results

### Disrupted peripheral T cell population homeostasis in *Rip1^tKO^* mice resembles that of aged mice

In accordance with the dynamic changes in T cells in aged humans^21–23^, aged mice (32 months old) experienced lymphopenia, with a decrease in T cells, particularly a 6% decrease in CD4+ T cells and a 4% decrease in CD8+ T cells in the spleen compared to young mice (3 months old) (**Figure 1A**). This decrease was especially notable in CD4 and CD8 naïve T cells, while memory T cells (CD44^high^) were predominant in the aged group (**Figure 1B**). Additionally, aged CD4 T cells differentiated towards T helper 1 (TH1) and regulatory T (Treg) subtypes, as the expression of master regulators T-bet and Foxp3 increased (**Figure 1C** and **D**). Furthermore, there was a significantly higher proportion of Gr1^+^CD11b^+^ cells (**Figure 1E**) and a noticeable thymus involution (**Figure 1F**) in the aged mice. The role of the immune system, especially T cells, in healthy aging emerges and become vital^24, 25^. In certain circumstances, T cells contribute to age-related diseases through the secretion of proinflammatory cytokines or the release of cytotoxic granules^26^, which suggests that T cell hyperactivation may directly cause the aging process.

**Fig 1.**
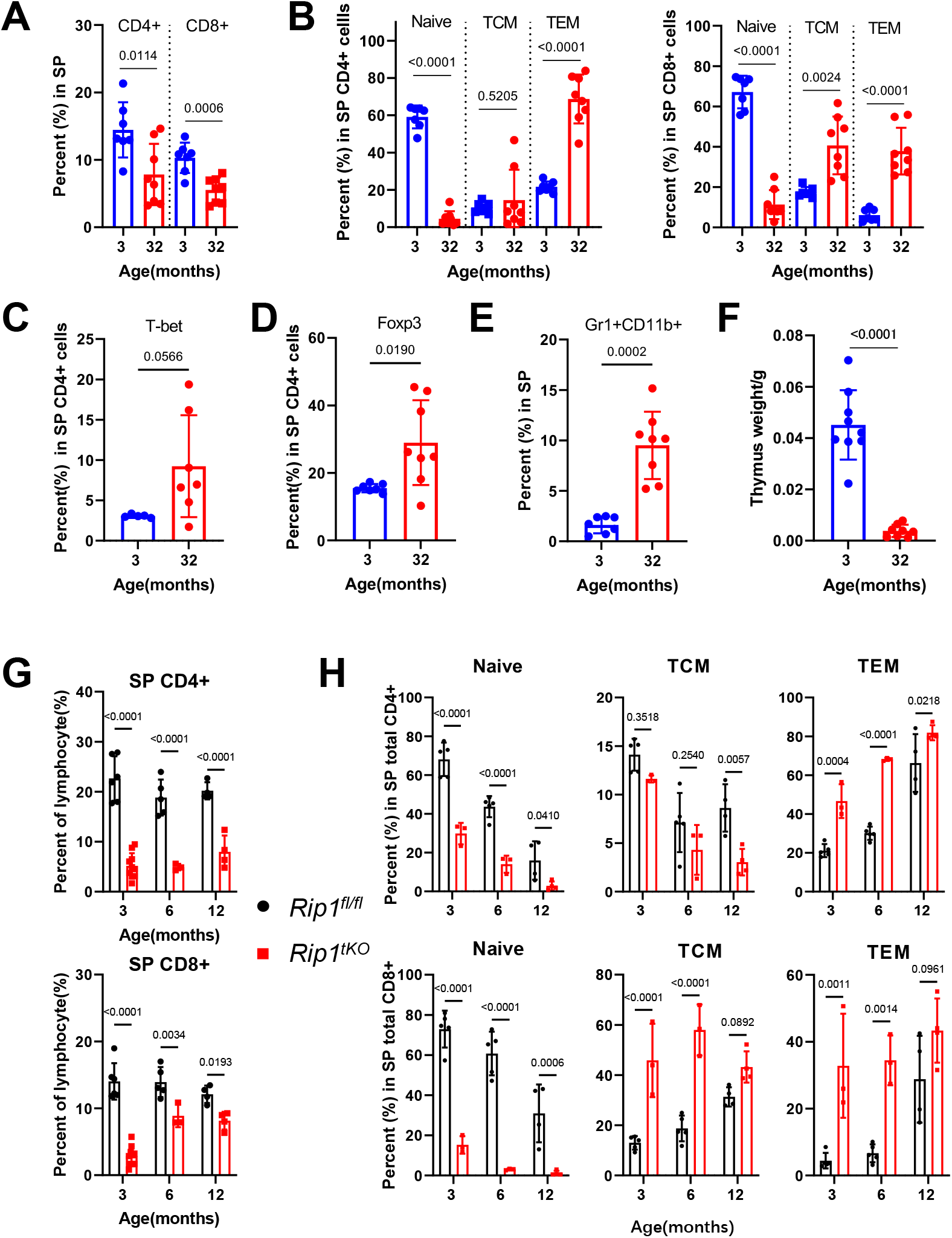
Disrupted peripheral T cell population homeostasis in *Rip1^tKO^* mice resembles aged mice. **A-D,** Flow cytometric quantification of splenic CD4^+^ and CD8^+^ T cell (A), T_naïve_ (CD44^-^CD62L^+^), TCM (CD44^+^CD62L^+^), TEM (CD44^+^CD62L^-^) (B) in 3-month-old and 32-month-old mice (n = 6 3-month-old; n = 8 32-month-old). Percentage of splenic CD4^+^ T cells positive for the Th1 cell transcription factor T-bet (C) and Treg cell transcription factor Foxp3 (D). **E,** Percentage of Gr1^+^CD11b^+^ cells in spleen (3-month-old and 32-month-old mice). **F,** Thymus weights of 3-month-old and 32-month-old mice. **G-H,** Flow cytometric quantification of splenic CD4^+^ and CD8^+^ T cell (H, n = 4 to 7), Tnaïve, TCM, TEM (I, n = 3 to 5) of the mice indicated at different age.

Given the key role of RIP1 in cell death and survival, a recent study has also reported that T cell-specific deletion of *Rip1* in mice leads to a noticeable decrease in peripheral T cell phenotype^20, 27^. This observation has prompted us to investigate whether mice lacking RIP1 in T cells can simulate the similar changes in T cell reduction seen in aged mice to study the relationship between T cell changes and individual aging. To determine this, we generated mice with a conditional knock-out of *Rip1* in T cells by crossing *Rip1^fl/fl^* mice with *Cd4-*Cre transgene mice that express the Cre recombinase under the control of the *Cd4* promoter^20^. Since the *Cd4-*Cre is expressed during the double positive period in the thymus, *Rip1* was deleted in all T cells (**Figure S1A**). Flow cytometric analysis revealed that the *Rip1^tKO^* mice had significantly fewer mature T cells (both CD4 and CD8) in the spleen compared to the aged-match control mice, regardless of their age (**Figure 1G** and **Figure S1B**). Naïve T cells in *Rip1^tKO^* mice were notably reduced at a young age and nearly disappeared by the time the mice reached 12 months old, while the population of memory and effector T cells increased (**Figure 1H** and **Figure S1C**). To confirm the changes in T cell pool observed in *Rip1^tKO^* mice, we crossed *Rip1^tKO^* mice with the *Rosa26^CAG-loxp-STOP-loxp-tdTomato^* (*R26^tdTomato^*) mice to generate T cells tracing mice (*Rip1^tKO^R26^tdTomato^*), in which all *Cd4*-Cre-expressing cells and their descendants are constitutively labeled by Tomato fluorescence. We also observed that the number of T cells was dramatically decreased in solid organs, including the kidney, liver, and heart in *Rip1^tKO^R26^tdTomato^* mice (**Figure S1D**). This demonstrates that RIP1 is critical in maintaining peripheral T cell homeostasis. The reduction of T cells and overactivation state in *Rip1^tKO^* mice resembles the T cell status of aged individuals, which prompted us to further investigate whether *Rip1*-deficient T cells could accelerate the aging process.

### The *Rip1^tKO^* mice displayed aggravated premature aging phenotypes and a shortened lifespan

Consistent with the previous report^20^, *Rip1^tKO^*mice appeared healthy and indistinguishable from *Rip1^fl/fl^* mice until adulthood. However, we observed physical deterioration in *Rip1^tKO^*mice around six months of age. Hindlimb clasping was first observed when *Rip1^tKO^*mice reached six months of age and became more severe over time, accompanied by other macroscopic changes such as hypotrichosis and greying of fur (**Figure 2A**). Both male and female *Rip1^tKO^* mice showed a tendency for resting weight gain (**Figure 2B** and **Figure S2A**). Upon further analysis at different ages, we observed severe thymus involution and a higher incidence of kyphosis in *Rip1^tKO^* mice (**Figure 2C, D** and **Figure S2B**), which are closely associated with advanced age.

**Fig 2.**
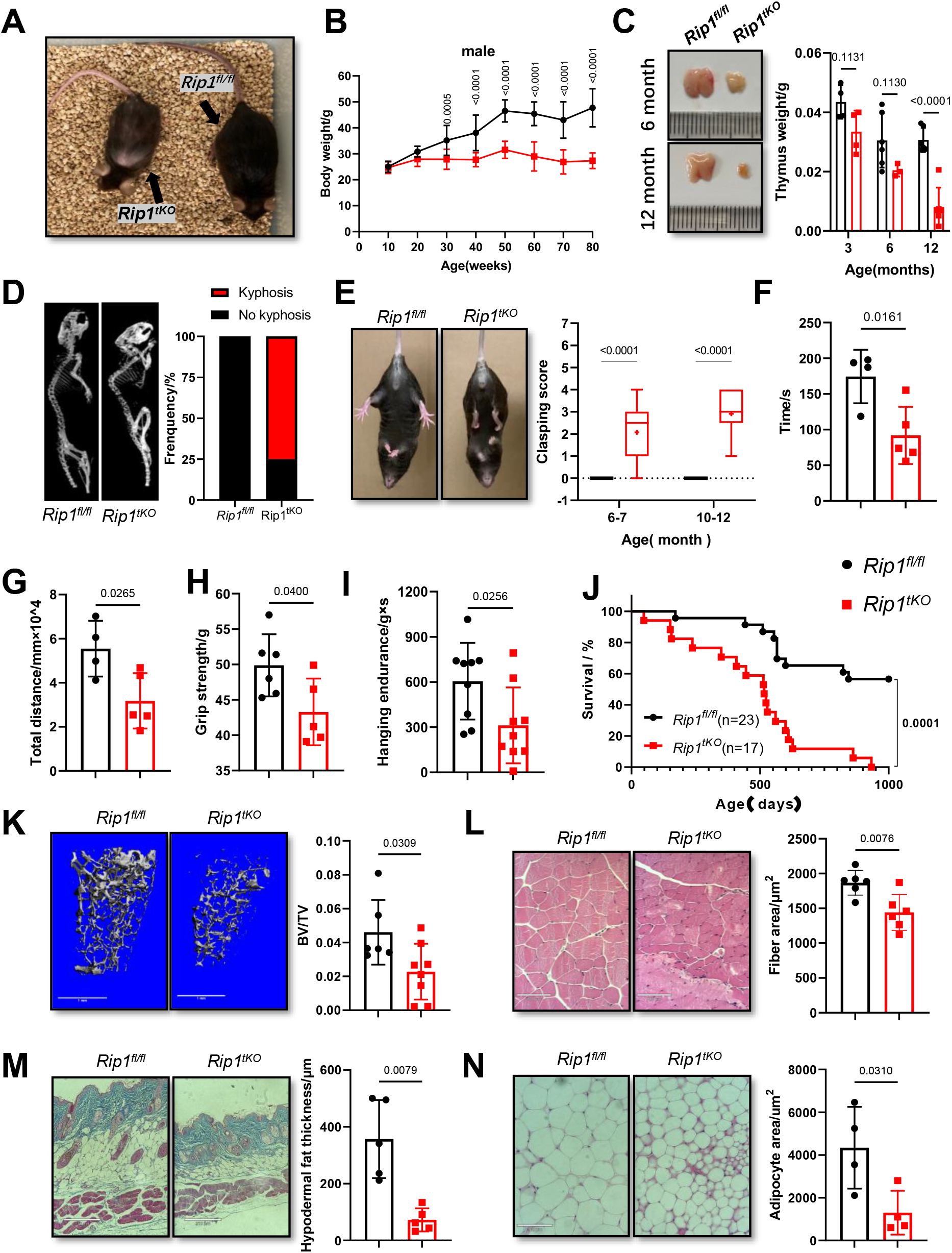
*Rip1^tKO^* mice develop premature aging phenotypes and age-related multi disfunctions. **A,** Representative photograph of 12-month-old female *Rip1^fl/fl^* and *Rip1^tKO^* mice. **B,** Body weight evolution in male *Rip1^fl/fl^* and *Rip1^tKO^* mice (n = 10 to 15). **C,** Representative photograph showing the thymic size of *Rip1^fl/fl^*and *Rip1^tKO^* mice at 6- and 12-month age (left). The graph (right) shows the thymus weight of *Rip1^fl/fl^* and *Rip1^tKO^*mice at 3-, 6- and 12-months age. **D,** Quantification of spine curvature by micro-computed tomography (CT) scans (left) and percentage of mice presenting lordokyphosis (right) in 12-month-old *Rip1^fl/fl^* and *Rip1^tKO^*mice (n = 10 to 11). **E,** Example of hindlimb clasping phenotype in 12-month-old *Rip1^fl/fl^*and *Rip1^tKO^* mice(left) and hindlimb clasping score (right; n = 10 to 19) in 20 seconds test at the age indicated. **F,** Rotarod performance by 12-month-old *Rip1^fl/fl^* and *Rip1^tKO^*mice, expressed as the average time spent on the rotating rod in four trials combined (n = 4 to 5). **G-I,** Total travel distance in open field test (G), grip strength (H) and hanging endurance (I) of 12-month-old *Rip1^fl/fl^*and *Rip1^tKO^* mice (n = 4 to 9). **J,** Kaplan-Meier survival curves for *Rip1^fl/fl^*and *Rip1^tKO^* mice (n = 14 to 23, including males and females). **K,** Representative Micro-CT 3D reconstructed images (left; scale bar, 1mm) and quantification of the femur trabecular bone volume fraction (BV/TV) (right) of 12-month-old male *Rip1^fl/fl^* and *Rip1^tKO^* (n = 5 to 8). **L,** Representative hematoxylin and eosin (H&E)–stained sections of the tibia anterior muscle (left; scale bar, 100μm) and quantification of mean myofiber cross-sectional area (right; n = 5 to 6; 10-month-old mice). **M,** Representative skin sections stained with Masson trichrome (left; scale bar, 210μm) and quantification of hypodermal fat thickness (right). The graph shows mean value of n = 5 mice at 12 months of age. **N,** Representative H&E-stained sections of gWAT (left; scale bar, 100μm) and quantification of mean estimated adipocyte surface area (right; n =4).

To assess the overall fitness of the animals, we tested their voluntary movement, coordination, muscle strength, and motor function. Notably, *Rip1^tKO^* mice showed higher clasping scores, poorer performance on the rotarod test, less movement in the open field test, decreased grip strength, and worse coordination on the hang-wire test (**Figure 2E** to **I** and **Figure S2C, D**), all of which ultimately led to a shortened lifespan (**Figure 2J**). Our observations suggest that *Rip1^tKO^* mice exhibited premature aging and eventually died prematurely.

To further evaluate the potential of *Rip1^tKO^* mice as an age-related disorders model, we carried out a comprehensive examination of the mice’s skeleton, muscle function, and lipid metabolism. We performed microcomputed Tomography (micro-CT) scans on the tibia and femur of 12-month-old mice, including both *Rip1^tKO^* mice and age-matched control mice. Our analysis of the 3D structural data indicated that the *Rip1^tKO^* mice developed osteoporosis, which was characterized by a loss of bone value in both the tibia and femur, a decrease in the number (Th.N) and thickness (Th.Th) of bone trabeculae, and an increase in trabecular space (Th.Sp) compared to the control mice (**Figure 2K**; **Figure S2E** and **F**). Additionally, histological analyses revealed a reduction in the mean fiber area of muscle tissue in the *Rip1^tKO^* mice compared to the control mice (**Figure 2L**). Another age-related pathology observed was thinning of the epidermis. The loss of subcutaneous fat in the *Rip1^tKO^*mice resulted in a significantly reduced skin thickness compared to the control mice (**Figure 2M**). *Rip1^tKO^* mice also had smaller adipocytes presented by histological analyses and loss of body fat mass measured by magnetic resonance imaging (MRI) (**Figure 2N** and **Figure S2G**). To summarize, our findings suggest that *Rip1* deficient in T cells causes osteoporosis, sarcopenia and lipolysis and results in premature aging and the progression of age-associated disorders.

### Progressive inflammation leads to senescence in distal tissues

Age-related disorders are usually characterized by chronic and progressive low-grade inflammatory^28–30^. To investigate the underlying mechanisms behind multimorbidity and frailty in the *Rip1^tKO^* mice, we performed primary T cell transcriptomics. The results of gene-ontology (GO) analysis showed that the upregulated differentially expressed genes in *Rip1*-deficient T cells were mainly enriched in inflammation and immune cell activation, such as activation of myeloid leukocytes and the interleukin-18 signaling pathway (**Figure 3A**). We also observed higher levels of circulating chemokines and cytokines (**Figure 3B** and **C**; **Figure S3A**) and an increased proportion of Gr1^+^CD11b^+^ in the spleen (**Figure 3D**) in 12-month-old *Rip1^tKO^* mice compared to that of age-matched control mice. Additionally, *Rip1*-deficient T cells secreted higher levels of the inflammatory cytokines IL-17A and interferon-γ (IFN-γ) (**Figure 3E** and **F**), the transcription factors T-bet and RORγt (**Figure 3G** and **Figure S3B**), as well as the transcription factor Foxp3 (**Figure S3C**) compared to WT T cells. These results demonstrated that *Rip1*-deficient T cells led to inflammaging by vibrant inflammatory cytokine production.

**Fig 3.**
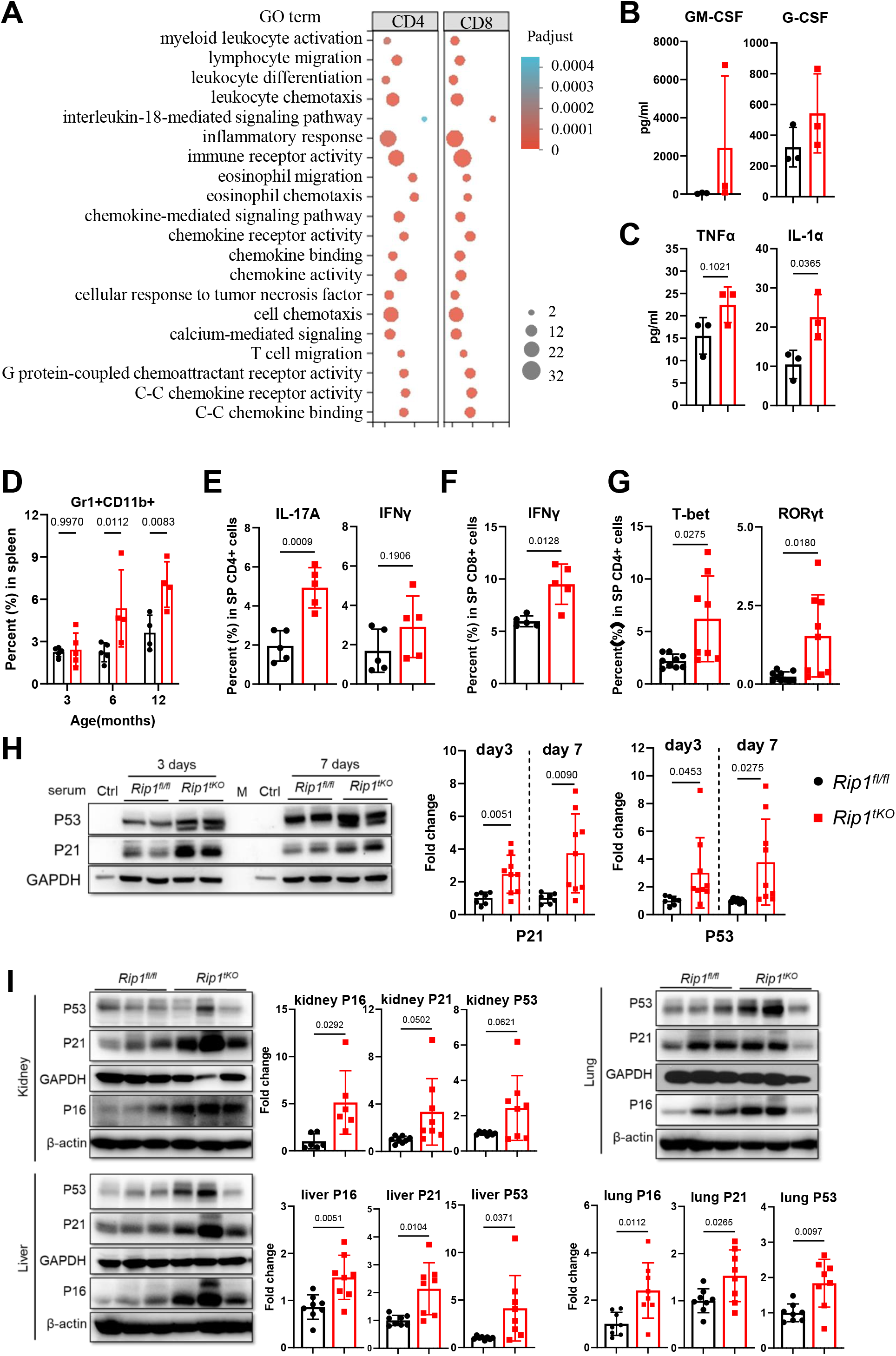
Progressive inflammation causes senescence in distal tissues. **A,** Gene Representative GO terms enriched in up-regulated, cytokine production-related DEGs, based on functional enrichment analysis in CD4^+^ and CD8^+^ T cells sorted from 2-month-old *Rip1^fl/fl^*and *Rip1^tKO^* mice. **B,** Serum levels of G-CSF and GM-CSF in 12-month-old *Rip1^fl/fl^* and *Rip1^tKO^*(n = 3). **C,** Percentage of Gr1^+^CD11b^+^ cells in spleens of *Rip1^fl/fl^* and *Rip1^tKO^* mice at indicated age (n = 4 to 5). **D-F,** Flow cytometric quantification of splenic CD4^+^ T cells staining positive for intracellular IL-17a and IFNγ (D), CD8^+^ IFN-γ^+^ T cells (E), Th1 cell transcription factor T-bet and Th17 cell transcription factor RORγt (F), (n = 5 to 9). **G,** Serum levels of inflammatory cytokines TNF-a and IL-1α detected by multiplex in 12-month-old *Rip1^fl/fl^*and *Rip1^tKO^* (n = 3). **H,** Representative immunoblot (left) and densitometry analysis (right) of P53 and P21 expression in primary WT mouse dermal fibroblasts (MDFs) cultured for 3 and 7 days in the presence of 12-month-old *Rip1^fl/fl^*or *Rip1^tKO^* mice serum. GADPH was used as loading control. Each lane represents independent mice serum used in the experiment. **I,** Representative immunoblot and quantification of P53, P21 and P16 protein expression in the lung, kidney, and liver from 12-month-old *Rip1^fl/fl^* and *Rip1^tKO^* mice. Loading controls were β-actin and GAPDH. Dots in all panels represent individual sample data.

Given that T cells may induce senescence through pro-inflammatory cytokines^26, 31^, we tested whether plasma pro-inflammatory cytokines from *Rip1^tKO^* mice could directly induce cellular senescence. We cultured non-senescent primary mouse dermal fibroblasts (MDFs) using medium mixed with 5% mouse plasma from 12-month-old mice. Compared to plasma from age-matched control mice, plasma from *Rip1^tKO^* mice increased the expression of the senescence genes P53 and P21^Waf/Cip1^ (**Figure 3H**). Additionally, *Rip1^tKO^* mice showed elevated protein levels of senescence markers in the heart, lung, kidney, liver, gonadal white adipose tissue, and intestine (**Figure 3I** and **Figure S3E**). Transcriptional level of p16 expression was also elevated in multiple organs in *Rip1^tKO^*mice (**Figure S3D**). We also detected an increase in senescence-associated β-galactosidase activity, a common biomarker for senescent cells in tissues and in culture^32^, in the kidney, lung, liver, and intestine of *Rip1^tKO^* mice (**Figure S3F**). These results suggest that *Rip1*-deficient T cells drive systemic senescence through their mediation of chronic inflammation.

### The apoptosis mediated by FADD plays a crucial role in maintaining T cell homeostasis in *Rip1^tKO^* mice

Since RIP1 serves as a key upstream regulator of necroptosis, apoptosis, and inflammation, and its death pattern is cell-type specific^33–38^, we aimed to uncover the downstream effector responsible for the observed lymphopenia and T cell hyperactivation in *Rip1^tKO^* mice. Given that RIP1 blocks both FADD/Capase-8-dependent apoptosis and RIP3/MLKL-dependent necroptosis during development^15, 16^, we crossbred *Rip1^tKO^* mice with *Rip3^-/-397^*, *Mlkl^-/-^40*, and *Fadd^-/-^Fadd:GFP^+^*^41, 42^ mice. The results showed that the absence of either RIP3 or MLKL had no impact on the reduced ratio and heightened activation of T cells in the spleen of *Rip1^tKO^* mice (**Figure 4A** and **B**). Furthermore, the hindlimb clasping, thinning of back skin (**Figure 4C**), and shortened lifespan (**Figure 4D**) remained unchanged in these mice, indicating that necroptosis is not involved in RIP1-regulated T cell homeostasis or age-related disorders in *Rip1^tKO^* mice.

**Fig 4.**
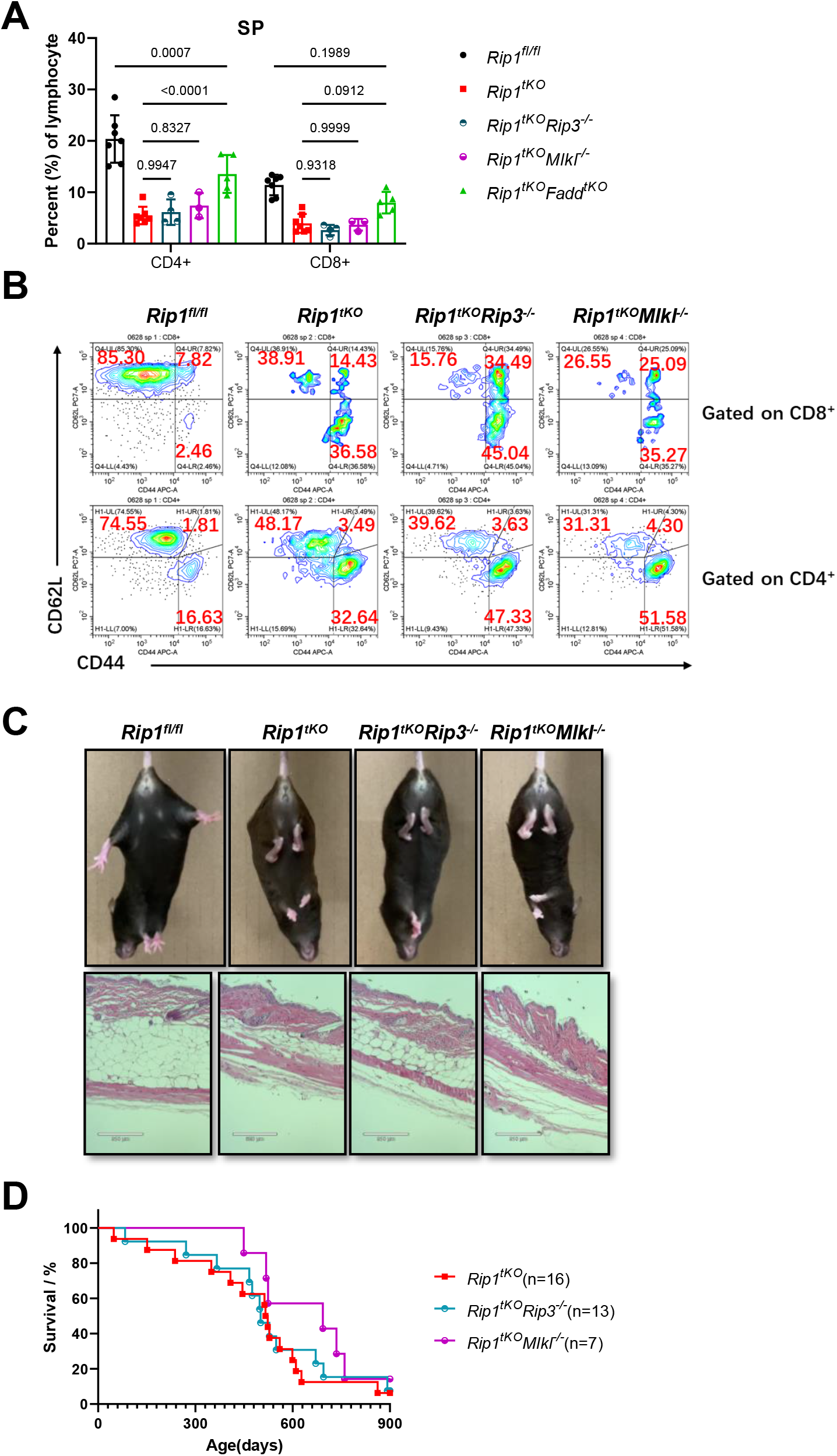
Blocking-up of necroptosis did not rescue T cell homeostasis and premature aging in *Rip1^tKO^* mice. **A,** Proportion of CD4^+^ and CD8^+^ T cells in *Rip1^fl/fl^*, *Rip1^tKO^*, *Rip1^tKO^Rip3^-/-^*, *Rip1^tKO^Mlkl^-/-^* and *Rip1^tKO^Fadd^tKO^* spleens determined by Flow cytometry (n=3 to 7; 8 weeks old). **B,** Flow cytometry of CD62L and CD44 expression in 8-week-old *Rip1^fl/fl^*, *Rip1^tKO^*, *Rip1^tKO^Rip3^-/-^*, *Rip1^tKO^Mlkl^-/-^* mice. **C,** Hindlimb clasping phenotype and skin HE staining images of the mice involved in (B) (mice were 10-12 months old). **D,** Kaplan-Meier survival curves for *Rip1^tKO^*, *Rip1^tKO^Rip3^-/-^*, *Rip1^tKO^Mlkl^-/-^* mice (n = 7 to 16, including males and females).

In contrast, eight-week-old *Rip1^fl/fl^Fadd^-/-^Fadd:GFP^+^Cd4^Cre^*mice (hereafter called *Rip1^tKO^Fadd^tKO^*) (**Figure S4A**) showed significantly recovered T cell percentage in spleen compared with *Rip1^tKO^* mice (**Figure 4A**), indicating the dominant role of apoptosis in T cell homeostasis. Additionally, we conducted T cell transcriptomics and observed that T cells sorted from *Rip1^tKO^* mice showed significantly up-regulated genes associated with apoptosis (**Figure 5A**), and the results were confirmed by western blot analysis, which revealed an increase in cleaved-caspase 3 expression levels in both CD4^+^ and CD8^+^ T cells from *Rip1^tKO^* mice compared to those of the control mice (**Figure 5B**). The proportion of CD4^+^ and CD8^+^ T cells in spleen was also restored significantly in *Rip1^tKO^Fadd^tKO^* mice at different age (**Figure 5C**). Importantly, in *Rip1^tKO^Fadd^tKO^* mice, naïve T cell pool was nearly completely recovered, and memory population shrank to the level comparable to control mice (**Figure 5D**). Furthermore, less Gr1^+^CD11b^+^ cells, Th1/Th17 T cell skewing and Treg cells were observed in the spleen of *Rip1^tKO^Fadd^tKO^*mice (**Figure 5E-G** and **Figure S4B**) compared to those of *Rip1^tKO^* mice, implying alleviated inflammation seen in the *Rip1^tKO^* mice. These results suggest that *Rip1*-deficient T cells died in an autonomous apoptosis manner which formed feedback, leading to T cell overactivation, which is restored by *Fadd* deletion in T cells. Thus, maintaining T cell homeostasis may be a critical element to sustain healthy physical condition.

**Fig 5.**
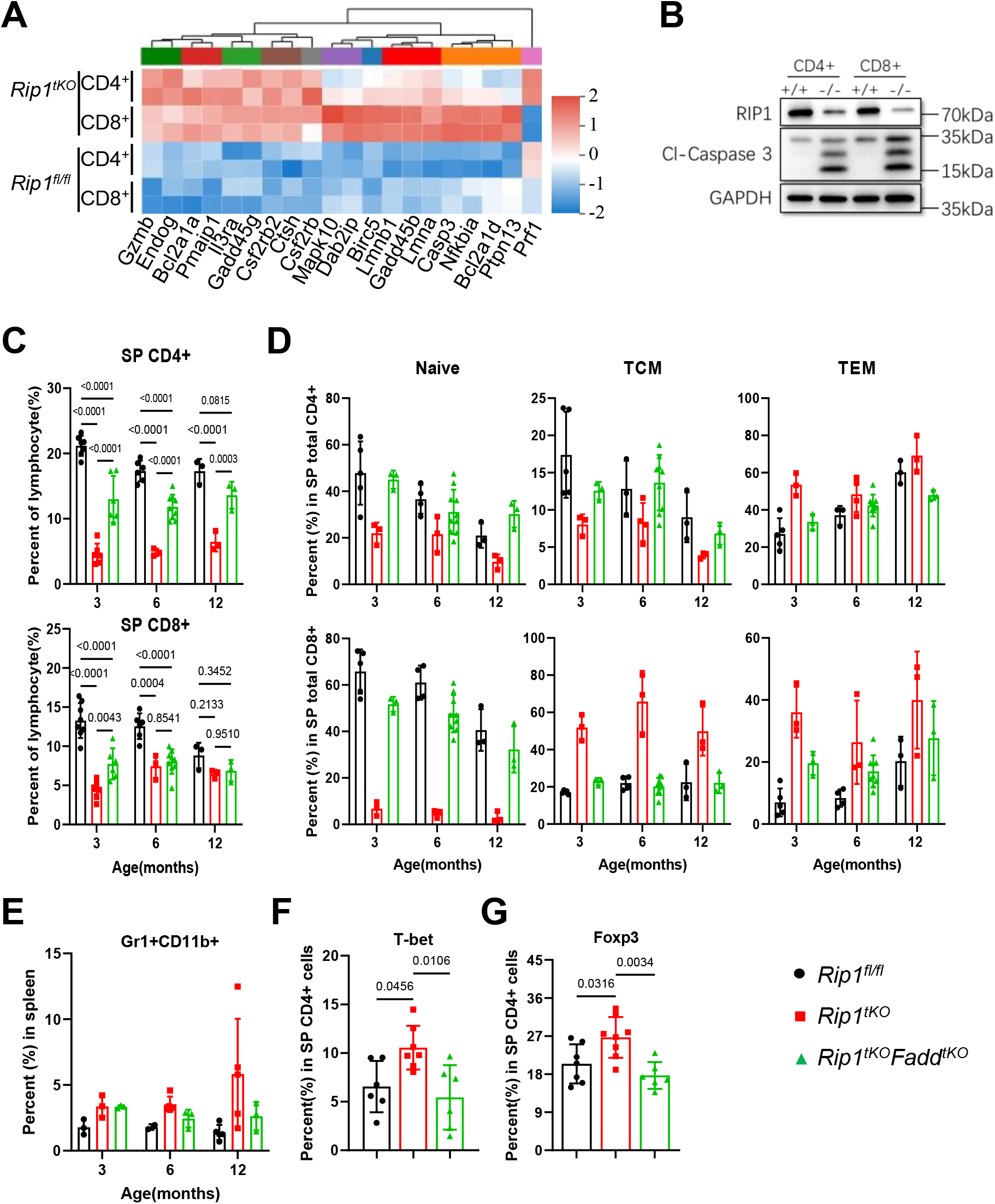
Blocking apoptosis in great part rescues peripheral T cell homeostasis in *Rip1^tKO^* mice. **A,** A heatmap depicting apoptosis-related genes were up regulated in *Rip1*-deficient CD4^+^ and CD8^+^ T cells. **B,** Representative immunoblot of apoptosis marker in CD4^+^ and CD8^+^ T cells sorted from 2-month-old mice (+/+ represent T cells isolated from *Rip1^fl/fl^* mice and -/- from *Rip1^tKO^*. Cl-caspase 3: cleaved-caspase 3. **C-D,** Flow cytometric quantification of splenic CD4^+^ and CD8^+^ T cell (C, n = 3 to 7), T naïve, TCM, TEM (D, n = 3 to 7) of *Rip1^fl/fl^*, *Rip1^tKO^* and *Rip1^tKO^Fadd^tKO^* mice at indicated age. **E,** Percentage of Gr1^+^CD11b^+^ cells in spleens of the mice at indicated age (n = 3 to 4). **F-G,** Percentage of splenic CD4^+^ T cells positive for T-bet (G) and Foxp3 (H) (n = 6 to 7).

### Blocking T cell apoptosis rescues premature aging and multimorbidity in *Rip1^tKO^* mice

Next, we analyzed the impact of deficiency of *Fadd* on premature aging and multimorbidity syndrome of *Rip1^tKO^* mice. Our results showed that *Rip1^tKO^Fadd^tKO^* mice had a longer lifespan (**Figure 6A**) with normal body weight gain (**Figure S4C**) and ameliorated thymus involution (**Figure 6B**) compared with *Rip1^tKO^* mice. Additionally, age-related disorders such as hindlimb clasping (**Figure 6C**), sarcopenia (**Figure 6E**), thinning of skin (**Figure 6G**), excessive lipolysis (**Figure 6H**), and osteoporosis (**Figure 6I**; **Figure S4E** and **F**) that were present in the *Rip1^tKO^* mice were fully prevented in the *Rip1^tKO^Fadd^tKO^*mice. Furthermore, blocking FADD-mediated apoptosis in T cells improved the physical quality of the *Rip1^tKO^* mice, as assessed by hanging endurance (**Figure 6D**), rotarod performance (**Figure 6F** and **Figure S4D**), and movement in open field test (**Figure S4G**). Finally, blocking T cell apoptosis prevented multiple instances of tissue senescence that were present in the *Rip1^tKO^* mice (**Figure 6J**; **Figure S4H** and **I**). These findings suggest that maintaining T cell homeostasis by blocking apoptosis in *Rip1^tKO^* mice can prevent premature aging and multimorbidity syndrome.

**Fig 6.**
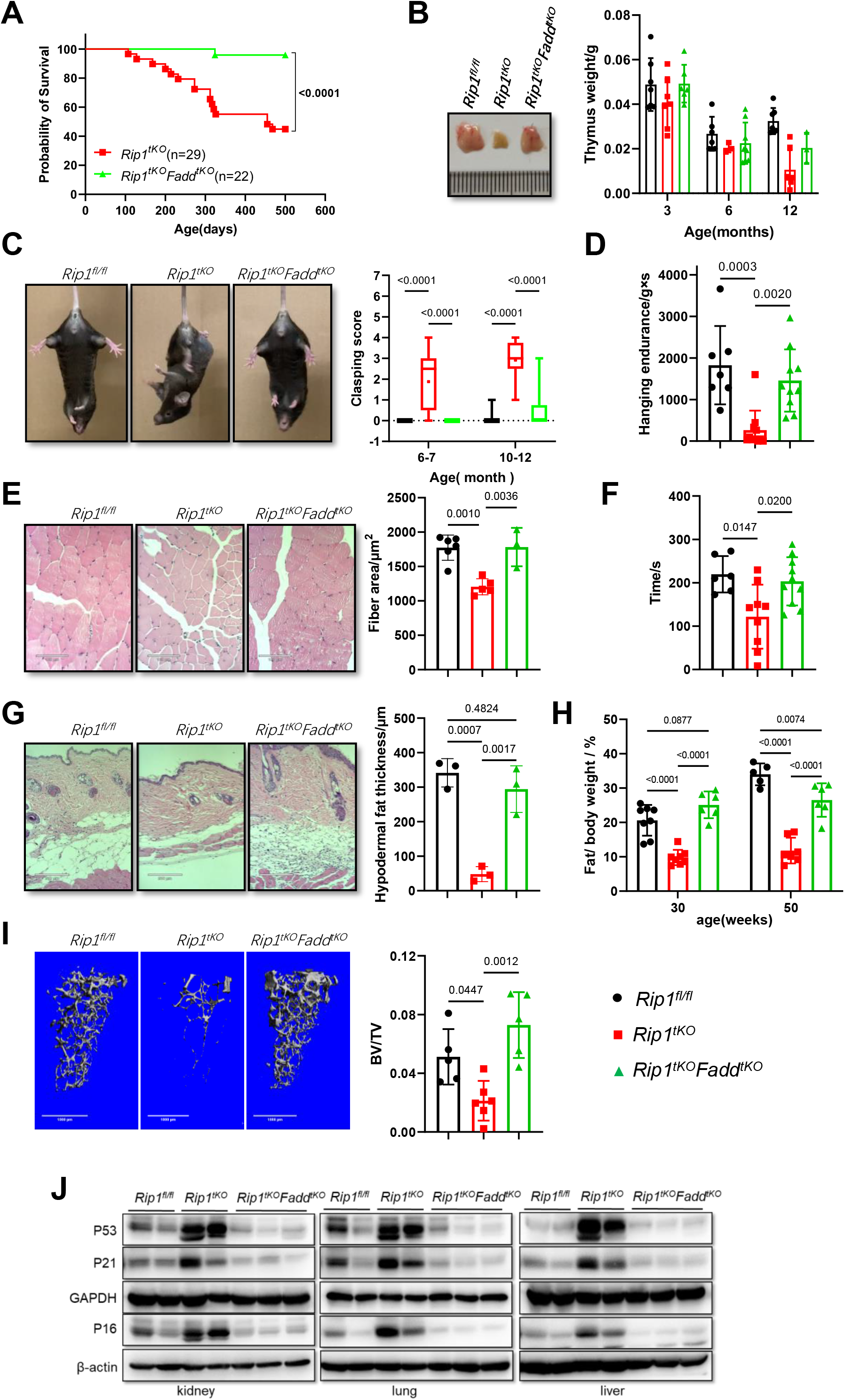
*Rip1^tKO^Fadd^tKO^* mice show lower rates of age-related disorders and counteract accelerated aging process. **A,** Kaplan-Meier survival curves for *Rip1^tKO^*and *Rip1^tKO^Fadd^tKO^*mice (n = 22 to 29, including males and females). **B,** Thymus photograph of *Rip1^fl/fl^*, *Rip1^tKO^* and *Rip1^tKO^Fadd^tKO^*mice at and 12-month age (left) and weights at 3-, 6- and 12-month age (n = 3 to 7). **C,** Hindlimb clasping phenotype in the 12-month-old mice (left) and hindlimb clasping score (right; n = 10 to 29) in 20 seconds test at the age indicated. **D,** Hanging endurance of the 12-month-old female mice (n = 7 to 10). **E,** Representative images of H&E-stained tibia anterior muscle from 12 months old mice (left; scale bar, 100μm) and quantification of mean myofiber cross-sectional area (right; n = 3 to 6). **F,** Rotarod performance by 12-month-old mice (n = 6 to 10). **G,** Representative H&E-stained skin sections (left; scale bar, 210μm) and quantification of hypodermal fat thickness (right) (n = 3; 12 months old mice). **H,** Adipose tissue percent determined by quantitative magnetic resonance imaging of mice at 30- and 50-week age (n = 6 to 8). **I,** Representative Micro-CT 3D reconstructed images (left; scale bar, 1mm) and quantification of the femur trabecular index BV/TV (right) of 12-month-old males (n = 5 to 6). **J,** Immunoblot of P53, P21 and P16 protein expression in the kidney, lung and liver from 12-month-old mice. Dots in all panels represent individual sample data.

It has been noticed that naïve T cells acquire memory phenotypes through homeostasis-driven proliferation^43, 44^, which is more active in individuals with lymphopenia^45^. Since *Rip1^tKO^* mice have a lymphopenic environment, it is highly possible that the increased T cell activation in these mice was due to homeostatic proliferation. To further investigate the mechanisms underlying T cell hyperactivation in *Rip1^tKO^* mice, we examined proliferation using Ki-67 staining. We found that a higher percentage of *Rip1*-deficient T cells, compared to wild-type cells, were positive for Ki-67 under normal conditions, indicating entry into the cell cycle, while regaining the normal lymphocytes number via blocking T cell apoptosis totally recovered the percentage of Ki-67 positive cells (**Figure S5A**). These results demonstrated that the absence of RIP1 in T cells, resulting in T cell hyperactivation, is due to lymphopenia-driven proliferation resulted from excessive apoptosis.

## Discussion

In this report, we demonstrate that specific T-cell *Rip1* deficiency mice experience age-related multimorbidity, including hair loss, hindlimb clasping, kyphosis, reduced motor coordination, osteoporosis, sarcopenia, abnormal fat metabolism, and ultimately premature death. This is due to excessive apoptosis in T cells, leading to compensatory proliferation and T cell hyperactivation, as well as increased levels of inflammation. Blocking necroptosis by crossing *Rip3^-/-^* or *Mlkl^-/-^* with *Rip1^tKO^* mice had no effect on aging phenotypes, while blocking apoptosis by deleting *Fadd* in T cells restored the balance between naive and memory T cells and rescued the aging-related phenotypes, ultimately extending lifespan of *Rip1^tKO^* mice. Our results suggest that the apoptosis of naive T cells is a driving factor in the aging process, and we proposed that replenishing or inhibiting apoptosis of naïve T cells has the potential to offer new therapies for aging and age-related diseases.

Recent years there is an explosion in the publication of research exploring the mechanisms underlying aging. In this study, we showed that shrinkage of the T cell pool and overactivation can cause premature aging and age-related multimorbidity, which is mediated by inflammaging in *Rip1^tKO^*mice. Additionally, *Rip1*-deficient T cells display a higher percentage of cells undergo cell cycle (as determined by Ki-67 staining) and acquire a highly differentiated and cytotoxic phenotype. Consistent with this, HIV-infected patients suffer from the death of CD4 T cells^46^ and subsequent chronic diseases, which resembles some aspects of immune senescence and premature aging^47–49^. Our findings emphasize the crucial role of maintaining normal T cell balance in preventing premature aging and provides a potential explanation for the many premature aging phenotypes observed in diseases such as AIDS, which is characterized by the decrease of T lymphocytes.

Previous study^50^ and our data, as shown in **Figure S6A**, indicate that the expression of RIP1 is significantly decreased in T cells obtained from aged individuals. T cells are constantly exposed to diverse antigens and undergo a continuous cycle of clonal expansion, death, and survival of memory subtypes^5^. However, the exact role of RIP1-regulated pathways in these processes is still unclear. We hypothesized that RIP1 plays a pro-survival role at certain stage of the T cell journey, while cell death serves as a checkpoint to control clonal expansion and to terminate uncontrolled responses. This could explain why the loss of RIP1 expression contributes to accelerated aging. Additionally, IL-7/IL-7R signaling is crucial for the homeostatic proliferation and survival of naive T cells^51, 52^, and the expression of IL-7R is impaired in peripheral T cells in *Rip1^tKO^*mice^27^. These findings further support the idea that RIP1 expression is necessary for the survival of peripheral T cells. Therefore, it is very necessary to further clarify the dynamic nature of RIP1 in T cells during aging.

The deletion of *Fadd* has a significant, although not complete, effect on restoring the T cell compartment in *Rip1^tKO^* mice. Interestingly, *Rip1^tKO^Fadd^tKO^* double knock-out mice appear even healthier than control mice, suggesting that there may be a minimum requirement for the number of T cells needed to maintain physiological and functional fitness. Our experiments obtained by necroptosis-blocking mice reveal that the RIP3/MLKL-mediated pathways are not involved in RIP1 knock-out T cells, demonstrating that the primary function of RIP1 in T cells is to protect against apoptosis. However, the specific molecular triggers that activate the death pathway are still veiled. Deletion of *Rip1* substantially increases the sensitivity of thymocytes to TNF-induced death^20^, so we crossed TNFR1 knock-out mice with *Rip1^tKO^* mice to test whether disrupting TNFR1-mediated signaling could rescue the peripheral T-cell defect. However, our results showed TNFR1-KO had little recovery effect on T cell quantity (**Figure S6B**), suggesting that either the TNFR2 pathway is still active or some other signaling molecules beyond TNFRs are involved.

Our study has uncovered that *Rip1^tKO^* mice exhibit signs of accelerated aging and premature death. By deleting the *Fadd* gene in *Rip1* deficient T cells, we observed a rescue of aging-related phenotypes and an extension of lifespan. In parallel investigation, Takayuki Imanishi et al. found that *Rip1* knock-out CD4 T cells showed higher activation levels of Akt, mTORC1, and ERK, leading to T cell senescence, and these phenotypes of T cells senescence can be restored by a combined deficiency of RIP3 and caspase-8^53^. These researchers concluded that the absence of RIP1 triggered the activation of RIP3/caspase-8 and mTORC1, contributing to T cell senescence and age-related diseases and premature death^53^. However, our genetic results showed that blocking necroptosis by deleting *Rip3* or *Mlkl* did not improve the premature aging and multiple diseases seen in *Rip1^tKO^* mice. In contrast, blocking apoptosis through the deletion of *Fadd* in T cells is enough to completely rescued the aging-related phenotypes and shortened lifespan in *Rip1^tKO^*mice. At this point, the specific mechanism by which *Rip1* deficiency in T cells accelerates aging is yet to be fully understood. Further studies are needed to shed light on the role of T cell aging-related changes in aging *in vivo*.

The mechanisms behind the decline of T cells with aging and its impact on various organs are not yet fully understood. T cells are essential for the immune system as they protect the body against pathogens and malignant cells. However, as the body ages, the quantity and effectiveness of T cells can decrease, resulting in a weakened immune response and increased susceptibility to infections and diseases^25^. Moreover, studies have indicated that T cells also play a role in regulating oxidative stress, a process where harmful byproducts of cellular metabolism can damage cells and tissues^26^. Despite this, the exact mechanisms behind how T cell aging-related changes trigger aging in various organs and eventually lead to death are still unclear. To gain a better understanding of these complex interactions between T cells and aging in different organs, the *Rip1^tKO^*mice model, which was induced by a specific factor causing T cell decline, can be used as a valuable tool.

Our findings have established that the decline of T-cells in *Rip1^tKO^*mice, resembles that of WT aged mice, which induces compensatory proliferation of T-cells leading to excessive activation, is one of the reasons for organ and individual aging and premature death. Restoration of the decrease in T-cells can restore organ and individual aging in *Rip1^tKO^* mice (**Figure S7**). These results suggest that it may be possible in the clinic to delay aging or treat age-related diseases by blocking excessive apoptosis of T-cells or by externally supplementing T-cells.

## Methods

### Mice

All mice used in this project were maintained in a specific pathogen-free (SPF) facility. *Rip1^fl/fl^* mice were generated by homologous recombination strategy (Shanghai Model Organisms Center, Inc.). In Brief, hybrid mouse ES cells were electroporated with a targeting vector containing floxed exon 3. The ES cells were selected by G418 and ganciclovoir, and screened for homologous recombination by PCR. Positive clones were then injected into C57BL/6 blastocysts, which were implanted into the uterus of pseudopregnant females to generate chimeras. These chimeras were then backcrossed with C57BL/6J mice for at least ten generations to generate *Rip1^fl/fl^*mice. *Cd4-*Cre transgene mice were kindly provided by Professor. Zichun Hua (Nanjing University). *Rosa26^CAG-loxp-STOP-loxp-tdTomato^*mice were obtained from Shanghai Model Organisms Center, Inc. *Rip3^-/-^*, *Mlkl^-/-^* and *Fadd^-/-^Fadd:GFP^+^* and *Tnfr1^-/-^* mice lines have been previously described^54–57^. All animal experiments were conducted in accordance with the guidelines of the Institutional Animal Care and Use Committee of the Institute of Nutrition and Health, Shanghai Institutes for Biological Sciences, University of Chinese Academy of Sciences.

### Reagents and antibodies

Antibodies used for western blotting listed as following: RIP1 (Cell Signaling Technology, 3493P), Cleaved caspase 3 (Cell Signaling Technology, 9661), FADD (Abcam, 124812), P53 (Cell Signaling Technology, 2524T), P21^Waf1/Cip1^ (BD Biosciences, 556431), P16^INK4a^ (Abcam, 211542), GADH (Sigma, G9545), β-actin (Sigma, A3854).

### Flow cytometry

Flow cytometric analysis was performed on Beckman CytoFlex S and data were analyzed with CytoExpert. After obtaining single cell suspensions, red blood cells were removed by incubation with a lysis buffer. For intracellular staining, Foxp3/Transcription Factor Staining Buffer Set (eBioscience, 00-5523-00) was used. For cytokine analysis, T cells were stimulated with PMA (Sigma, P1585) and ionomycin (Sigma, I3909) for 4 hours followed brefeldin A stimulation for another 2 hours. All the samples in the same experiments and comparisons were gated under the same parameters.

Sorting of T cells were performed on the Beckman Moflo Astrios sorter. Mouse total primary T cells were isolated from the spleen and peripheral lymph nodes of mice by EasySep Mouse T Cell Isolation (Stemcell, 19851) and then were further sorted by flow cytometric cell sorting based on CD3^+^CD4^+^ and CD3^+^CD8^+^ surface markers. The purified cells were used according to experimental design.

The following antibodies were used: against mouse CD45 (BD, 553080), CD3e (eBioscience, 25-0031-82; BD, 553061), CD4 (eBioscience, 47-0042-82), CD8e (Biolegend, 100732 and 100738), B220 (eBioscience, 17-0452-83), Gr-1 (eBioscience, 12-5931-83), CD11b (eBioscience, 17-0112-83), CD62L (eBioscience, 25-0621-81), CD44 (eBioscience, 17-0441-81), T-bet (Invitrogen, 25-5825-80), RORγt (Invitrogen, 12-6988-80), GATA3 (Biolegend, 653805), Foxp3 (eBioscience, 17-0441-81), IFNγ (Biolegend, 505825), IL-17A (Biolegend, 506919), Ki67 (Biolegend, 652403), LIVE/DEAD (Biolegend, 423101).

### Western blot

Cell pellet or tissue were lysed in RIPA lysis buffer containing 1× proteinase inhibitor cocktail (Roche, 5056489001), 2 mM PMSF and 1 mM DTT. After centrifugation, the protein concentration in the collected supernatants was determined using a BCA protein assay kit (Thermo Scientific, 23225). The normalized lysates were subsequently denatured in reducing 5x SDS loading sample buffer at 100 °C for 5 min.

### Micro-CT scanning and Magnetic resonance imaging (MRI)

Micro-CT scanning was used for kyphosis and osteoporosis determination. Angles below 15% of the average controls were classified as kyphosis^8^. Mice were deeply sedated using isoflurane (RWD, R510-22-8) during image acquisition. Regions of interest (ROI) were drawn for tibia and femur. Body fat content measurement was assessed by MRI (EchoMRI).

### Hindlimb clasping score

For this test, mice were suspended gently by the tail and videotaped for 10-15 seconds. Three separate trials were taken for each mouse. Hindlimb clasping severity was rated from 0 to 4:

0 = hindlimbs splayed outward and away from the abdomen,

1 = one hindlimb retracted inwards towards the abdomen intermittently,

2 = both hindlimbs partially retracted inwards towards the abdomen intermittently,

3 = both hindlimbs continuously retracted inwards towards the abdomen.

4 = one or both hindlimbs continuously retracted inwards towards the abdomen plus foot weakness while walking across a wire cage top.

Scores of 0.5 were utilized when appropriate.

### Physical function assessments

For hanging test, mice were place onto the metal cage top, which was then inverted and suspended above the home cage; the latency to when the animal falls is recorded. Three days of practice was performed before official test. This test is performed three days per week with three trials per session. The average performance for each session is presented as the average of the three trials.

For open field test, mice were placed in the box for 20 mins and total travel distance, the time and track in center zone or edge zone were recorded using Cleversys software. For rotarod test, mice were trained on the rotarod first. On the test day, mice were placed onto the rotarod, which was started at 4 r/m and accelerated from 4 to 40 r/m during a 5-min-test. The latency to when the animal falls from rotarod is recorded. Results averaged from four trails over a 30-min interval.

For grip strength (g) test, a Grip Strength Meter (Columbus Instruments) was used, with results averaged over 15 trials. To measure grip strength, the mouse is swung gently by the tail so that its forelimbs contact the bar. The mouse instinctively grips the bar and is pulled horizontally backwards, exerting tension. When the tension becomes too great, the mouse releases the bar.

### SA β-galactosidase assay

SA-β-gal staining was performed using kits from Beyotime (C0602) and Cell Signaling Technology (9860S), according to the manufacturer’s instructions. Briefly, OCT-embedded tissue sections on glass coverslips were washed in PBS, fixed and washed, then incubated with SA-β-Gal staining solution overnight at 37℃ following the manufacturer’s instructions. After completion of SA-β-Gal staining, sections were stained with nuclear fast red (Servicebio, G1035) for 5-10 min at room temperature, rinsed under running water for 1 min. Sections were dehydrated in increasing concentrations of alcohol (70%-80%-95%-100% -100%) and cleared in xylene twice. After drying, samples were examined under a bright-field microscope.

### MDF culture

Primary mouse dermal fibroblasts (MDFs) were separated from the skin of newborn mice. For senescence induction by mouse serum, MDFs were cultured in Opti-MEM medium (Gibco, 31985070) with 5% serum from 12-month-old *Rip1^fl/fl^*and *Rip1^tKO^* mice, 1% penicillin, and 100 µg/ml streptomycin for 3 or 7 days, then cell lysis was collected for further analysis.

### ELISA and Luminex detection of proinflammatory cytokines and chemokines

Serum cytokines and chemokines were detected with Bio-Plex Pro Mouse Cytokine Grp I Panel 23-plex (#M60009RDPD) using the Bio-Plex 200 system (Luminex Corporation, Austin, TX, USA). TNF-α (eBioscience, 88-7324-88), IL-1β (eBioscience, 88-7013-88), IL-6 (eBioscience, 88-7064-88), IFN-β (Biolegend, 439407) ELISA kits were used as well.

### RNA extraction and Real-Time PCR

Total RNA was extracted from frozen tissues with Trizol reagent (Takara, 9108) and was retrotranscribed to cDNA with PrimeScript™ RT reagent Kit with gDNA Eraser (Takara, RR047A). Real-time quantitative RT-PCR (qPCR) reactions were performed with TB Green Premix Ex Taq™ (Takara, RR420A) in 384-well plates in a QuantStudio™ 6 Flex Real-Time PCR System. The following primers were used for qPCR: P16 (5’-CGTACCCCGATTCAGGTG-3’, 5’-ACCAGCGTGTCCAGGAAG-3’), β-actin (5’-CTAAGGCCAACCGTGAAAAG-3’, 5’-ACCAGAGGCATACAGGGACA-3’), GAPDH (5’-TGTGTCCGTCGTGGATCTGA-3’, 5’-CCTGCTTCACCACCTTCTTGAT-3’). Relative expression was calculated as RQ = 2^−ΔΔCt^.

### RNASeq

Total RNA was extracted from CD4 and CD8 T cells sorted from 8-week-old mice and was measured using the NanoDrop2000 (Thermo Fisher Scientific), agarose gel electrophoresis and Agilent2100. RNA samples displayed a 260/280 ratio of around 2.0 were used. The cDNA libraries were constructed and sequenced by Majorbio Biotech. In brief, 1ug RNA for each sample was used for cDNA libraries construction by Illumina TruseqTM RNA sample prep Kit . The constructed DNA was enriched by PCR amplification for 15 cycles and then purified by 2% gel electrophoresis. Clone clusters were generated on the Illumina cBot and high-throughput sequencing was performed on Illumina Novaseq 6000 sequencer using NovaSeq Reagent Kits v1.5 and 2 × 150 bp paired-end reads were generated. The data were analyzed on the online tool of Majorbio Cloud Platform (https://cloud.majorbio.com/page/tools/).

### Statistical analysis

Image-Pro Plus 6 was used in image and immunoblot quantification. Data in figures are presented as mean ± s.d., and statistical P value was determined by unpaired two tailed Student’s t-test (comparing two groups), one-way ANOVA or two-way ANOVA (comparing more than two groups), log-rank test with GraphPad Prism 9 primed.

## SUPPLEMENTARY INFORMATION

Supplementary information includes six figures.

### AUTHOR CONTRIBUTIONS

L.X.W and H.B.Z designed the study; L.X.W carried out experiments and analyses with assistance from M.L, X.M.L, Y.J.O, X.M.Z, X.X.W, X.H.W, J.L.L, M.Y.X, H.L, Y.Z, Y.C.T, F.L, J.S.D and X.J.Z; X.X.Z and H.W.Z conducted the analysis of mouse phenotypes and K.L.L assisted with the histopathological analyses.; J.B.L and Y.W.Z. provided technical support; X.Z.W, Y.L, B.Z provided resources, intellectual input and edited the manuscript; L.X.W and H.B.Z assembled figure panels and wrote the paper. H.B.Z supervised the project.

## ACKNOWLEDGEMENTS

We would like to thank Dr. Xiaodong Wang (National Institute of Biological Sciences, Beijing, China) for providing *Ripk3^-/-^*mice, Dr. Jianke Zhang (Thomas Jefferson University, Philadelphia, PA, USA) for providing *Fadd^-/-^Fadd:GFP^+^* mice and Dr. Feng Shao (National Institute of Biological Sciences, Beijing, China) for providing *Tnfr1^-/-^*mice. We also thank Lin Qiu (Shanghai Institute of Nutrition and Health, Chinese Academy of Sciences) for technical assistance and Zhonghui Weng (Shanghai Institute of Nutrition and Health, Chinese Academy of Sciences) for mice maintained. This work was supported by grants from National Key Research and Development Program of China (2022YFA0807300), the Strategic Priority Research Program of Chinese Academy of Sciences (XDA26040306, XDB39000000), National Natural Science Foundation of China (32270803, 31970688, 82272181). Shanghai Excellent Academic/Technical Leader Program (22XD1404500) and Shanghai Science and Technology Commission (23141902800). We also thank support from Shanghai Frontiers Science Center of Cellular Homeostasis and Human Diseases, and GuangCi Professorship Program of Ruijin Hospital Shanghai Jiao Tong University School of Medicine.

## CONFLIC OF INTEREST

The authors declare no competing interests.

---

**Supplementary Figure 1.**
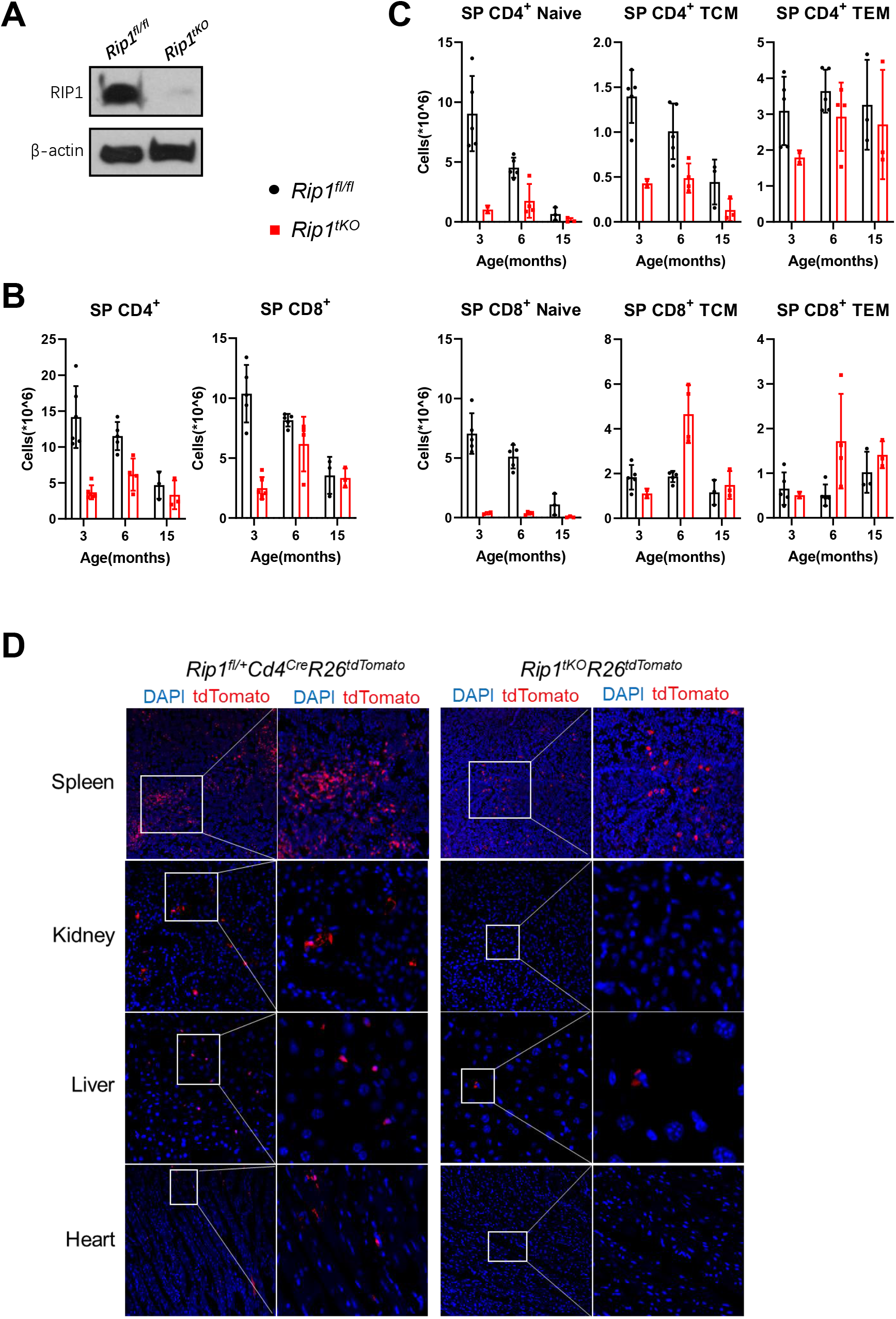
Disrupted peripheral T cell population homeostasis in *Rip1^tKO^* mice. **A,** Immunoblot detection of RIP1 in total T cell (CD3^+^) isolated from peripheral lymph node and spleen of 2-month-old *Rip1^fl/fl^* and *Rip1^tKO^* mice. **B-C,** Absolute number of splenic CD4^+^ and CD8^+^ T cell (B, n = 4 to 7), T naïve, TCM, TEM (C, n = 3 to 5) of the mice indicated at different age. **D,** Representative immunofluorescence (IF) images of *ROSA26^+^* cells in spleen, kidney, liver, and heart in *Rip1^tKO^R26^tdTomato^*and *Rip1^fl/+^Cd4^Cre^R26^tdTomato^*mice.

**Supplementary Figure 2.**
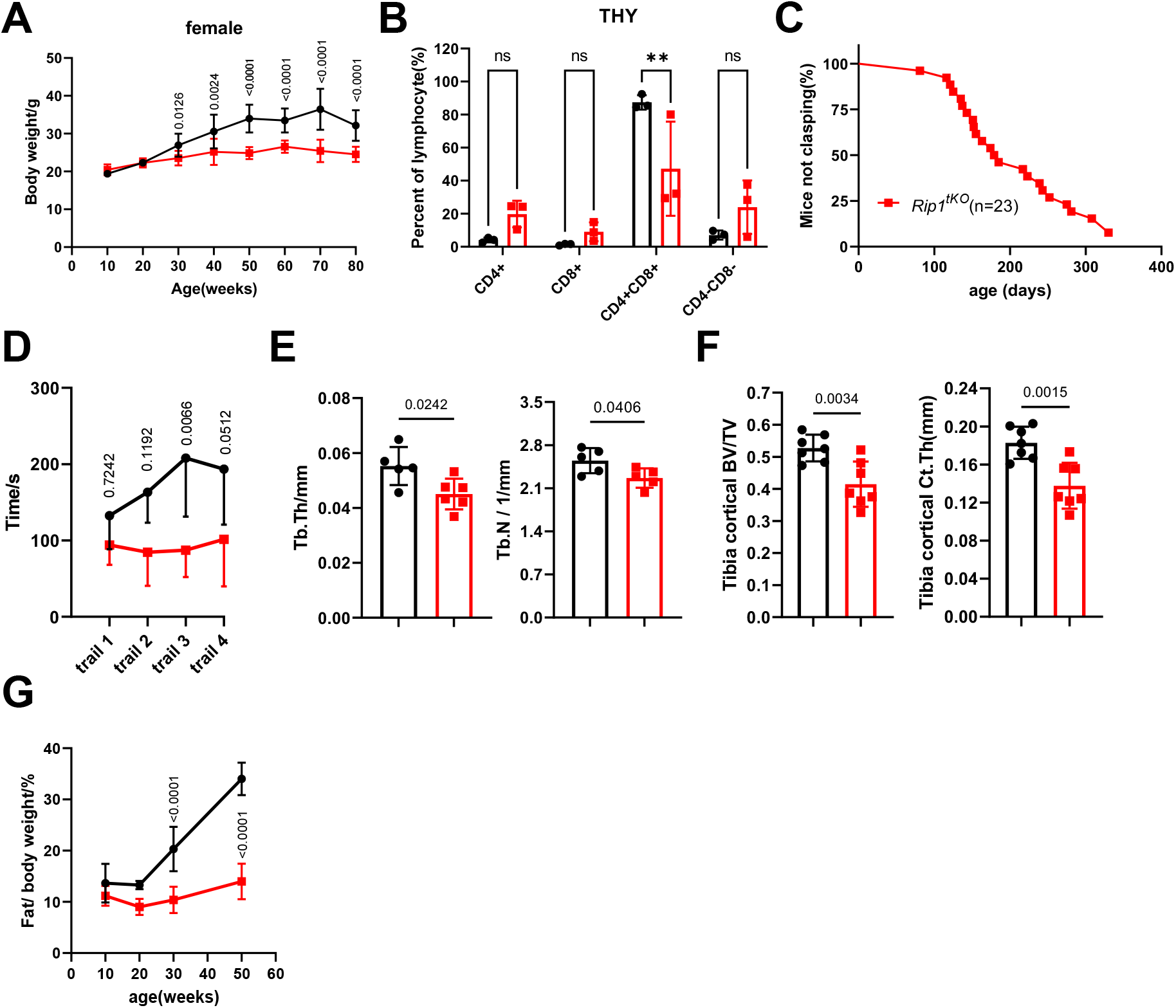
Premature aging phenotype of *Rip1^tKO^* mice. **A,** Body weight evolution in female *Rip1^fl/fl^* and *Rip1^tKO^* mice (n = 10 to 15). **B,** Flow cytometric quantification of CD4^+^, CD8^+^, CD4^-^CD8^-^ and CD4^+^CD8^+^ subtype in thymus, mice at age of 12 months (n=3). **C,** Percent of and *Rip1^tKO^* mice not observed hind climb clasping during adult life. **D,** Rotarod performance by 12-month-old *Rip1^fl/fl^* and *Rip1^tKO^*mice, expressed as the time spent on the rotating rod in four separate trials (n = 4 to 5). **E,** Micro-CT scan quantification of the femur trabecular index of 12-month-old male *Rip1^fl/fl^* and *Rip1^tKO^* mice (n = 5 to 8). Trabecular thickness (Tb.Th), trabecular number (Tb.N). **F,** Micro-CT quantification of tibia cortical bone index of 12-month-old male *Rip1^fl/fl^* and *Rip1^tKO^* mice (n = 7). **G,** Adipose tissue percent determined by quantitative magnetic resonance imaging of mice at 10-, 20-, 30- and 50-week age (n = 4 to 12). **H,** Time course (left) and average oxygen expenditure (right) during 24-hour cycle obtained using metabolic cages (n = 6 to 7). Mice were 10-month-old.

**Supplementary Figure 3.**
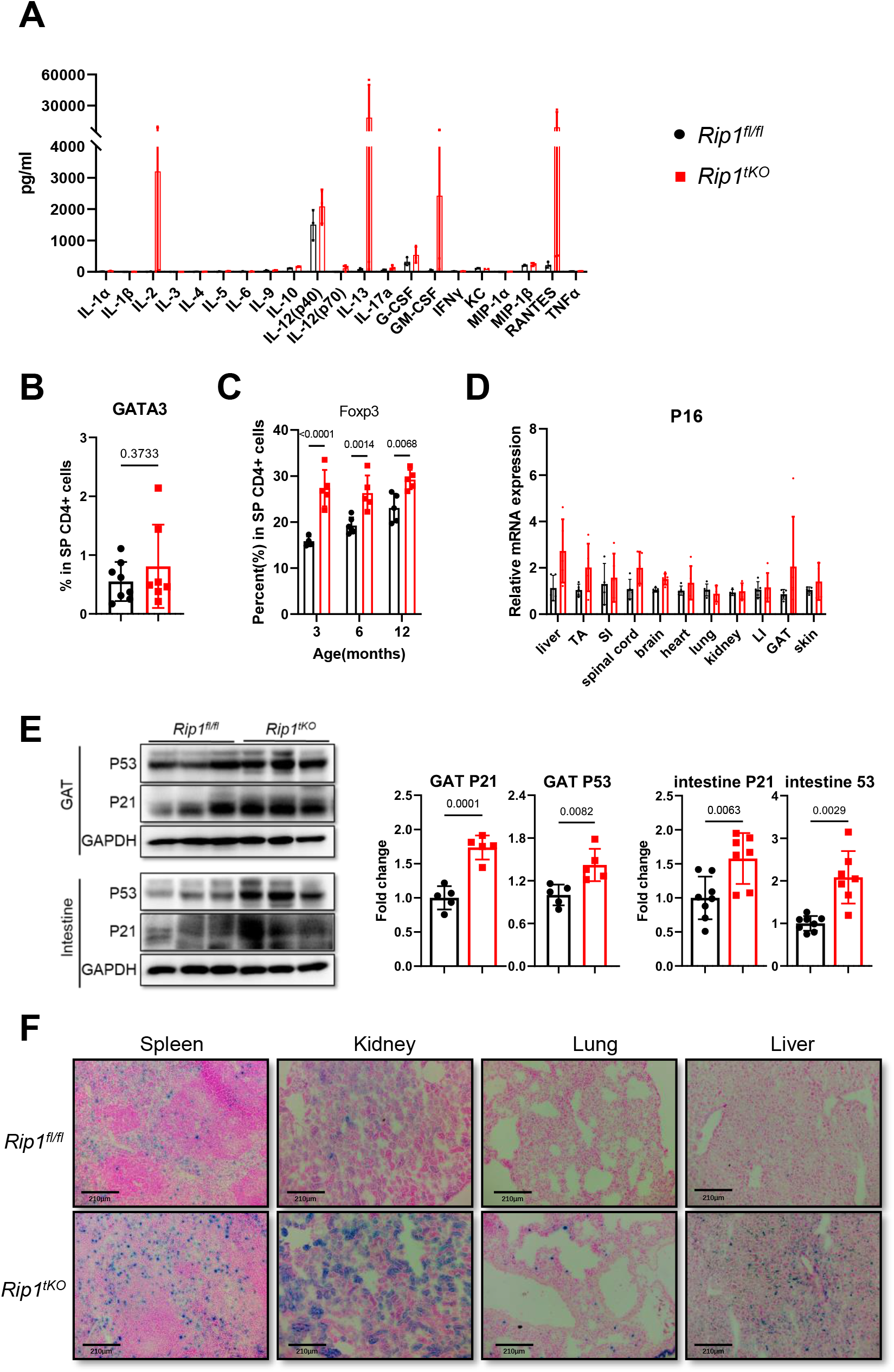
Inflammaging induces premature senescence in *Rip1^tKO^* mice. **A,** Serum levels of cytokines and chemokines assayed by Luminex liquid suspension chip detection (n=3; 12 months old). **B,** Percentages of GATA3 positive cells in total splenic CD4^+^ T cells (n=7 to 8; 3-month-old mice). **C,** Percentages of Foxp3 positive cells in total mice splenic CD4^+^ T cells at the age indicated (n = 3 to 6). **D,** Relative mRNA expression of P16 in multi organs of 12-month-old mice (n=5 to 6). Representative images of senescence-associated β-galactosidase (SA-β-galactosidase) staining of spleens, kidneys, lungs, and livers from 12-month-old *Rip1^fl/fl^* and *Rip1^tKO^* mice (scale bar, 210μm). **E,** Representative immunoblot (left) and protein fold change quantification (right) for P21 and P53 for GAT and intestine (n = 5 to 7; 12-month-old mice). Dots in all panels represent individual sample data. **F,** Representative senescence-associated β-galactosidase (SA-β-galactosidase) staining of spleens, kidneys lungs and livers from 12-month-old mice (scale bar,210μm).

**Supplementary Figure 4.**
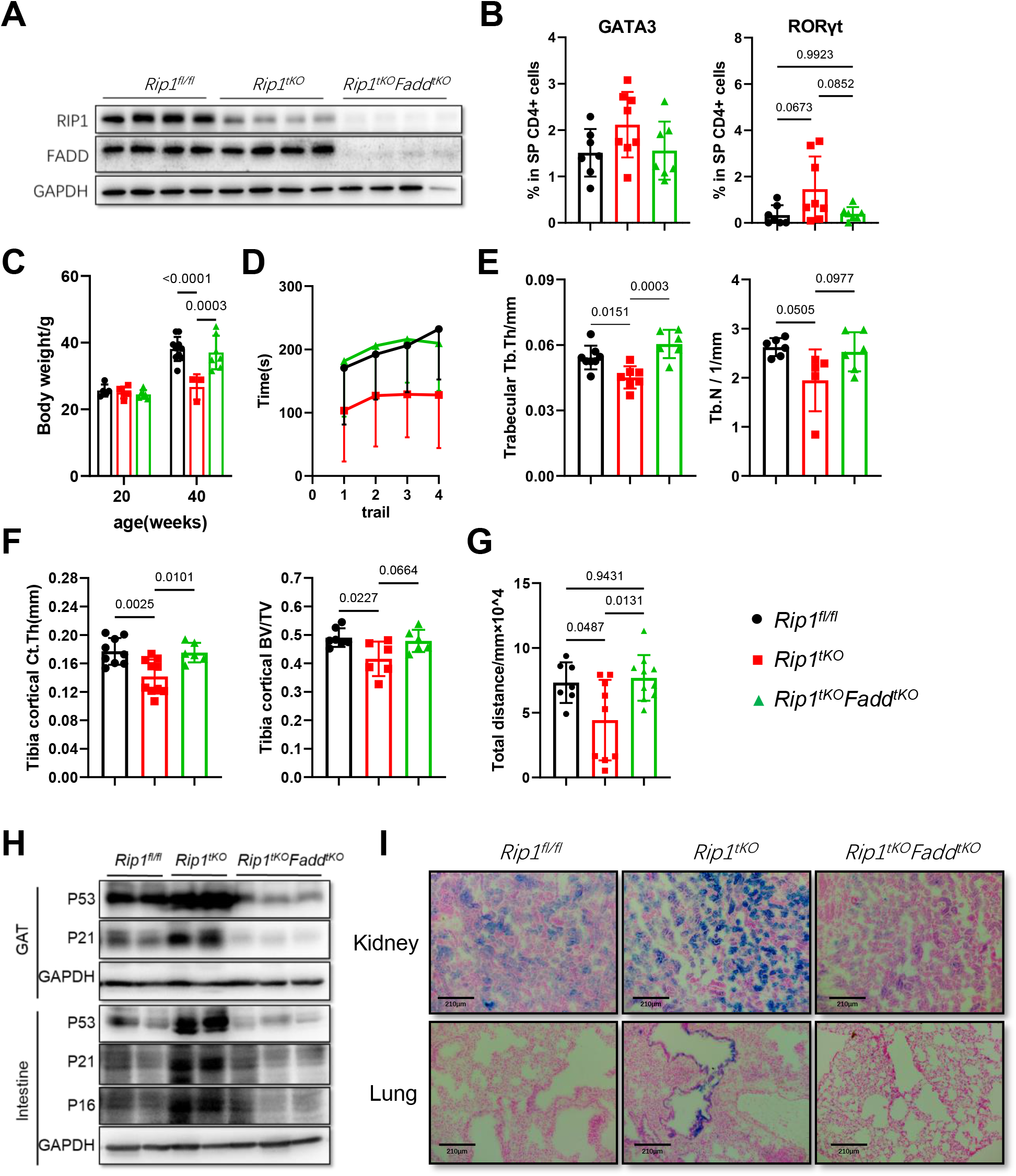
*Rip1^tKO^Fadd^tKO^* mice showed attenuated premature aging symptoms. **A,** Immunoblot detection of RIP1 and FADD in total T cell isolated from peripheral lymph node and spleen of 2-month-old *Rip1^fl/fl^*, *Rip1^tKO^* and *Rip1^tKO^Fadd^tKO^* mice. **B,** Percentages of GATA3 and RORγt positive cells in total splenic CD4^+^ T cells (n = 7 to 8). **C,** Body weight of mice at 10- and 40-week-old age (n = 3 to 7, male). **D,** Rotarod performance in four separate trials by 12-month-old mice (n = 6 to 10). **E,** Micro-CT scan quantification of the femur trabecular index Tb.Th and Tb.N of 12-month-old males (n = 5 to 6). **F,** Quantification of tibia cortical bone index of 12-month-old male (n = 6 to 7). **G,** Total travel distance in open field test by 12-month-old mice (n = 7 to 10) **H,** Immunoblot for P16, P21 and P53 for GAT and intestine from 12-month-old mice. Each lane represents individual sample data. **I,** Senescence-associated β-galactosidase (SA-β-galactosidase) staining of kidneys and lungs from 12-month-old mice (scale bar,210μm).

**Supplementary Figure 5.**
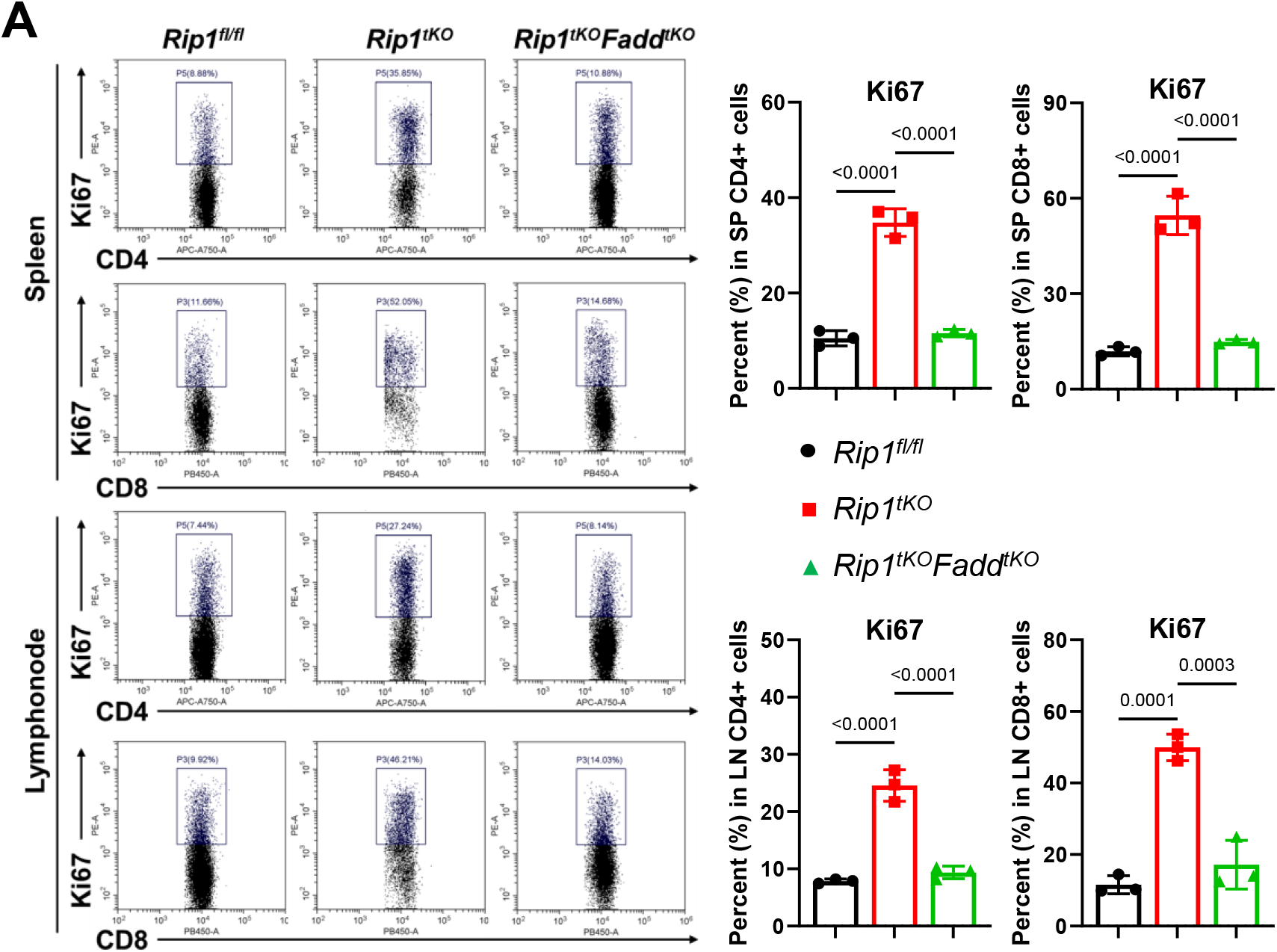
Loss of quiescence and hyperactivation of RIP1 deficient T cells is an apoptosis-dependent manner. **A,** Flow cytometry and quantification of Ki-67 positive CD4 or CD8 T cells from spleen and lymph node. Mice were 8-week-old.

**Supplementary Figure 6.**
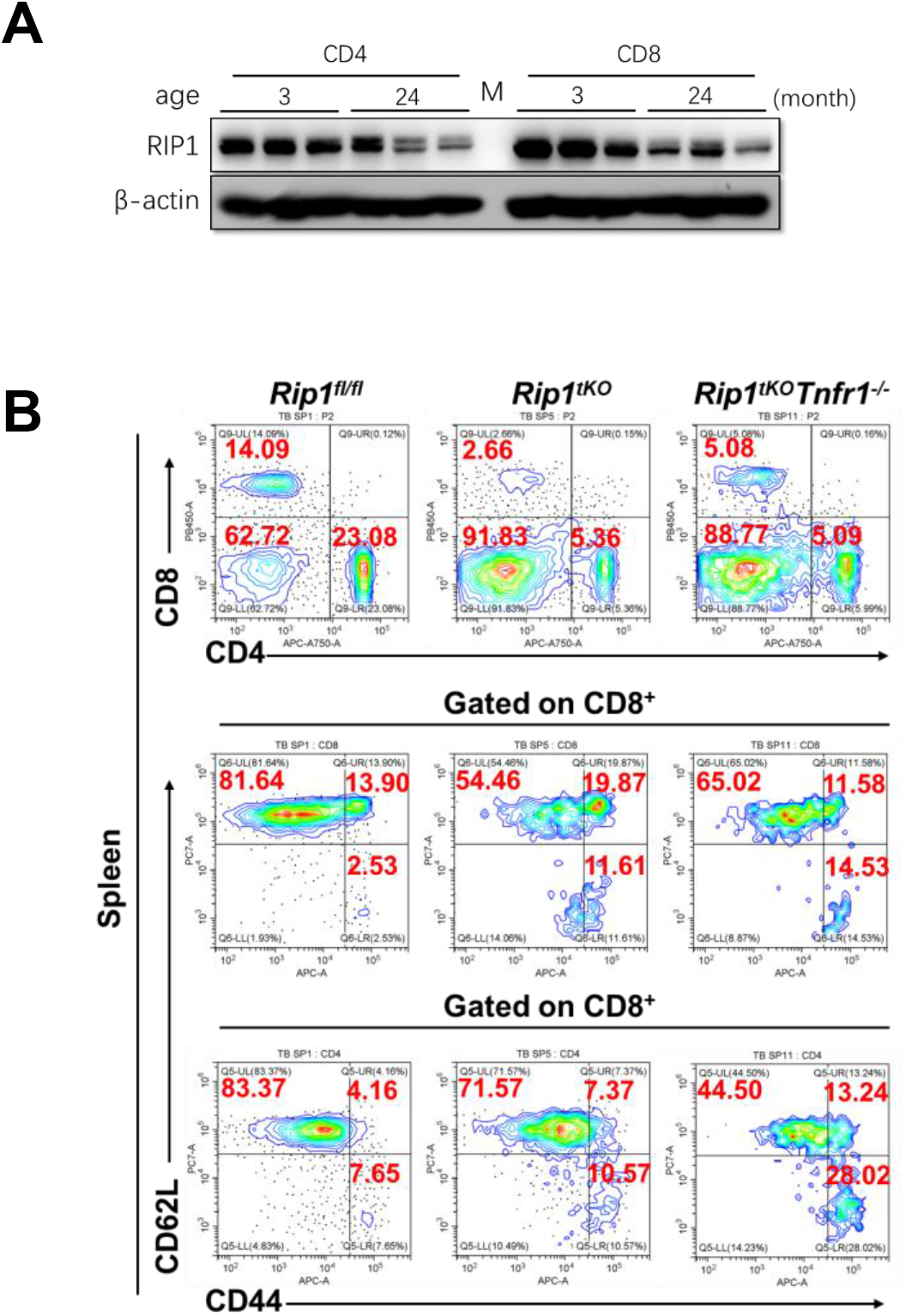
**A,** Immunoblot of RIP1 expression in CD4^+^ and CD8^+^ T cells sorted from spleen of 3- and 24-month-old wild type mice. Each lane represents individual sample data. **B,** Representative dot plots of CD4^+^ and CD8^+^ (top) T cells, and naïve/effector/ memory cells T cells from the spleens of 6-week-old *Rip1^fl/fl^*, *Rip1^tKO^* and *Ript^tKO^Tnfr1^-/-^*mice.

**Supplementary Figure 7.**
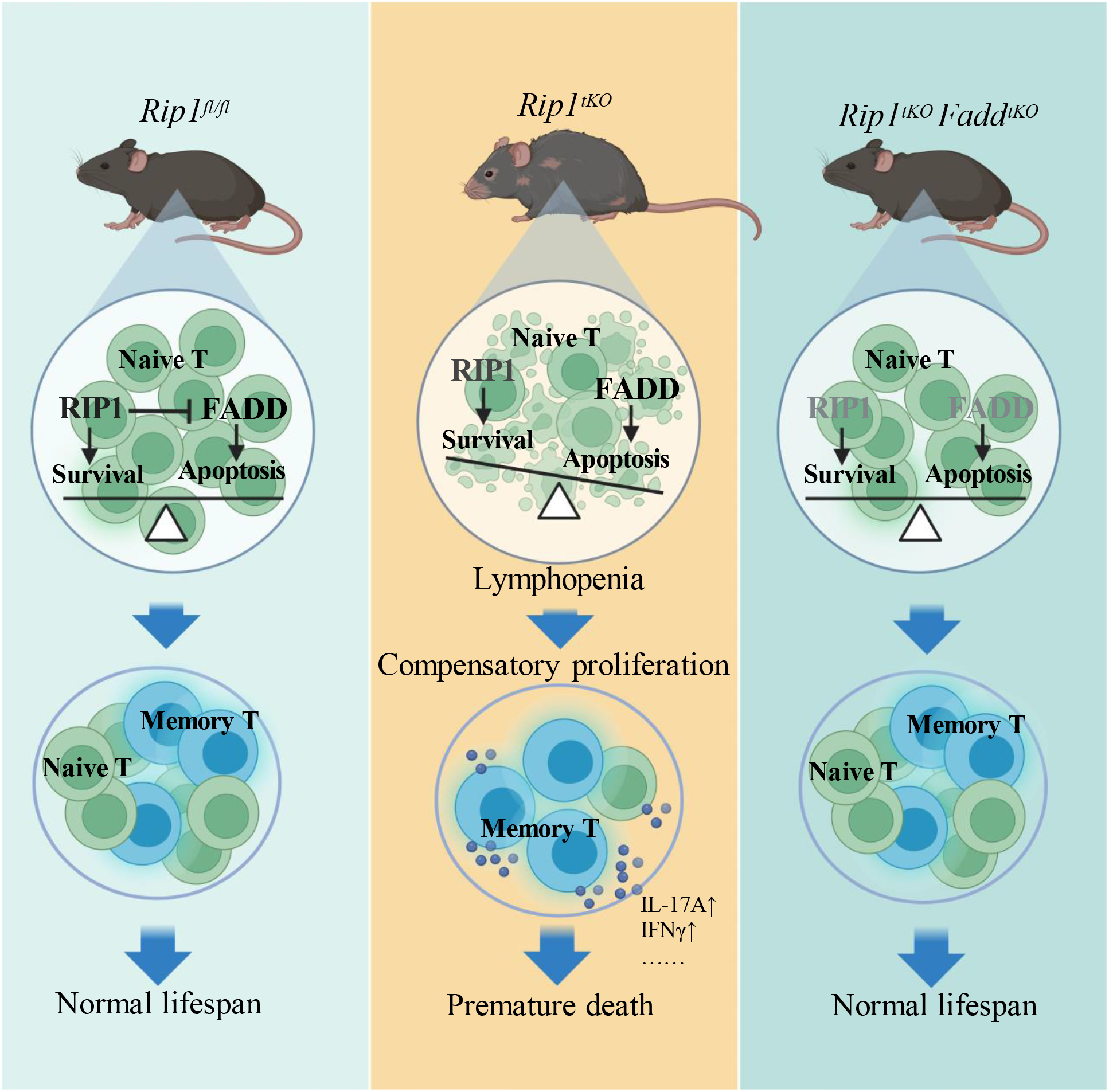
Model for premature death caused by *Rip1* deficient T cells. In *Rip1^fl/fl^* mice, RIP1 inhibits FADD mediated apoptosis in T cells, ensuring the T cell homeostasis. In *Rip1^tKO^* mice, T cells lacking RIP1 exhibit excessive FADD mediated apoptosis, resulted in lymphopenia driven compensatory proliferation, which prompts T cell hyperactivation with increased pro-inflammatory cytokine secretion, ultimately leading to premature death of *Rip1^tKO^* mice. In *Rip1^tKO^Fadd^tKO^*mice, T cells apoptosis is blocked by *Fadd* deletion, which restores the T cell homeostasis, leading to a normal life span compared to *Rip1^tKO^*mice.

